# Direct PA-binding by Chm7 is required for nuclear envelope surveillance at herniations

**DOI:** 10.1101/2020.05.04.074880

**Authors:** David J. Thaller, Danqing Tong, Christopher J. Marklew, Sapan Borah, Barbara Ciani, C. Patrick Lusk

## Abstract

Mechanisms that control nuclear membrane remodeling are essential to maintain the integrity of the nucleus but remain to be fully defined. Here, we identify a phosphatidic acid (PA)-binding activity in the nuclear envelope-specific ESCRT, Chm7, in budding yeast. PA-binding is mediated through a conserved hydrophobic stretch of amino acids, which confers specific binding to the inner nuclear membrane (INM). This INM-binding is independent but nonetheless required for interaction with the LAP2-emerin-MAN1 (LEM) domain protein, Heh1 (LEM2). Consistent with the functional importance of PA-binding, mutation of this region inhibits recruitment of Chm7 to the INM and abolishes Chm7 function in the context of nuclear envelope herniations or “blebs” that form during defective nuclear pore complex (NPC) biogenesis. In fact, we show that PA accumulates at nuclear envelope herniations. We suggest that local control of PA metabolism is important for ensuring productive nuclear envelope remodeling and that its dysregulation may contribute to pathologies associated with defective NPC assembly.

## Introduction

The nuclear envelope provides a selective barrier that ensures the integrity of nuclear-cytoplasmic compartmentalization. It is now clear that disruption of nuclear envelope function, either by direct perturbation of the nuclear membranes, or by inhibition of the assembly of nuclear pore complexes (NPCs), is determinantal to cell viability, can result in losses in genome integrity, and can cause disease (Jevtić and Levy, 2014; Hatch and Hetzer, 2014; Lusk and King, 2017; Ungricht and Kutay, 2017; Shah et al., 2017; Houthaeve et al., 2018). Perhaps not surprisingly, cells have evolved protective mechanisms that surveil and ameliorate disruptions of the nuclear compartment, including defective *de novo* NPC biogenesis (Thaller and Lusk, 2018).

NPC biogenesis during interphase is thought to occur through an inside-out mechanism that begins at the inner nuclear membrane (INM)(Otsuka et al., 2016). As the NPC is built from ∼30 proteins (termed nucleoporins or nups) radially constructed in multiples of 8, hundreds of nups are ultimately assembled to build a single NPC (Kosinski et al., 2016; Kim et al., 2018; Allegretti et al., 2020). Although the molecular “steps” in NPC assembly remain obscure, a key event is the fusion of the INM and outer nuclear membrane (ONM) to form a nuclear pore. The underlying molecular mechanism that drives INM-ONM fusion is unknown (Rothballer and Kutay, 2013; Weberruss and Antonin, 2016; Hampoelz et al., 2019). Genetic evidence is mounting, however, that local changes in lipid metabolism may contribute to the NPC biogenesis mechanism, likely at the step of INM-ONM fusion (Schneiter et al., 1996; Siniossoglou et al., 1998; Scarcelli et al., 2007; Hodge et al., 2010; Lone et al., 2015; Zhang et al., 2018). For example, the formation of nuclear envelope blebs or herniations that appear over malformed NPCs might occur due to an inhibition of INM-ONM fusion (Wente and Blobel, 1993; Scarcelli et al., 2007; Onischenko et al., 2017; Zhang et al., 2018; Thaller et al., 2019; Allegretti et al., 2020; Rampello et al., 2020). These structures have been observed in both yeast and metazoan systems (Thaller and Lusk, 2018) and have been associated with defects in lipid metabolism (Schneiter et al., 1996; Grillet et al., 2016).

Nuclear envelope herniations associated with NPC misassembly may also be caused by triggering a nuclear envelope surveillance pathway that monitors the integrity of nuclear-cytoplasmic compartmentalization (Webster et al., 2014, 2016; Thaller et al., 2019). The principal components of this pathway include the endosomal sorting complexes required for transport (ESCRT) and integral INM proteins of the LAP2-emerin-MAN1 (LEM) family including Heh1 in budding yeast (a.k.a Src1; LEM2 in higher eukaryotes)(Webster et al., 2014, 2016). In recent work, we showed that the ESCRT Chm7 (orthologue of CHMP7) has nuclear export sequences (NESs) that restrict its access to the nucleus and Heh1 (Thaller et al., 2019). CHMP7 also has an NES (Vietri et al., 2019). Perturbations to the nuclear transport system or nuclear membranes is predicted to disrupt the spatial segregation of Chm7 and Heh1, which leads to their binding and the subsequent activation of an ESCRT-dependent nuclear envelope repair pathway (Thaller et al., 2019; Lusk and Ader, 2020). The molecular mechanisms of nuclear envelope repair remain obscure, but as ESCRTs also seal the nuclear envelope at the end of an open mitosis (Olmos et al., 2015; Vietri et al., 2015; Gu et al., 2017; Ventimiglia et al., 2018; Willan et al., 2019; Warecki et al., 2020; Pieper et al., 2020; Gatta et al., 2020), it seems likely that a Chm7-Heh1 partnership leads to the closure of nuclear pores through an ill-defined annular fusion event. As the hyper-activation of this pathway with gain-of-function mutants of Chm7 leads to aberrant membrane remodeling and expansion at the INM (Thaller et al., 2019, Vietri et al., 2019), there is an interesting relationship between changes in local lipid synthesis and membrane repair that remains to be mined.

Indeed, recent work in *C. elegans* has uncovered synthetic genetic interactions between ESCRT genes (including CHMP7) and regulators of phosphatidic acid (PA) metabolism (Penfield et al., 2020). As PA is a major precursor of both phospholipids and triacylglycerol, this pathway decides whether to generate new phospholipids for membrane growth or to store fatty acids in lipid droplets (Carman and Han, 2011), some of which have been shown to grow from the INM (Romanauska and Köhler, 2018). Further, PA can have a potent destabilizing effect on membrane structure (Kooijman et al., 2003; Kwolek et al., 2015), which can be deleterious but can also be leveraged to promote membrane fusion reactions (Corrotte et al., 2006; Nakanishi et al., 2006; Zeniou-Meyer et al., 2007; Liu et al., 2007; Yang et al., 2008; Giridharan et al., 2013; Bates et al., 2014; Adachi et al., 2016; Starr et al., 2016; Miner et al., 2017; Park et al., 2019), often by PA-binding amphipathic helices in key membrane remodeling proteins (Chernomordik and Kozlov, 2003; Zhukovsky et al., 2019). As such, PA levels are kept under tight control; whether changes in local PA can directly impact nuclear envelope remodeling events remains unknown.

In our continuing efforts to determine the mechanism of nuclear envelope surveillance, we discovered a specific PA-binding activity in Chm7. This interaction is important for Chm7’s function as mutation of a likely amphipathic helix disables its ability to be recruited and activated at the nuclear envelope. Through these efforts, we uncovered evidence that nuclear envelope herniations associated with NPC misassembly likely contain high levels of PA. Thus, local control of PA at the nuclear envelope is a key feature of essential nuclear envelope remodeling events.

## Results and Discussion

### A conserved hydrophobic stretch and NES2 are required for Chm7 function

We, and others, have previously identified several functional domains within Chm7 that directly contribute to Chm7 localization at steady state, and when nuclear envelope surveillance is triggered (Olmos et al., 2016; Webster et al., 2016; Gu et al., 2017; Vietri et al., 2019; Thaller et al., 2019; Gatta et al., 2020; von Appen et al., 2020). For example, we had identified two sequence elements in the C-terminal ESCRT-III domain that are sufficient to act as NESs (Fig. 1 A; also see (Thaller et al., 2019)). It remained unclear, however, whether one or both of these sequences were necessary to prevent accumulation of Chm7 in the nucleus. Similarly, we had not yet investigated a role for an interesting hydrophobic stretch (“H”, Fig. 1 A) that was first identified in the N-terminus of the human protein (Olmos et al., 2016) and appears to be conserved, although less hydrophobic, in budding yeast (Fig. S1 A, B). Thus, to fill in these gaps in our understanding of Chm7, we generated point mutants in each of these motifs and assessed their localization and function.

**Figure 1.**
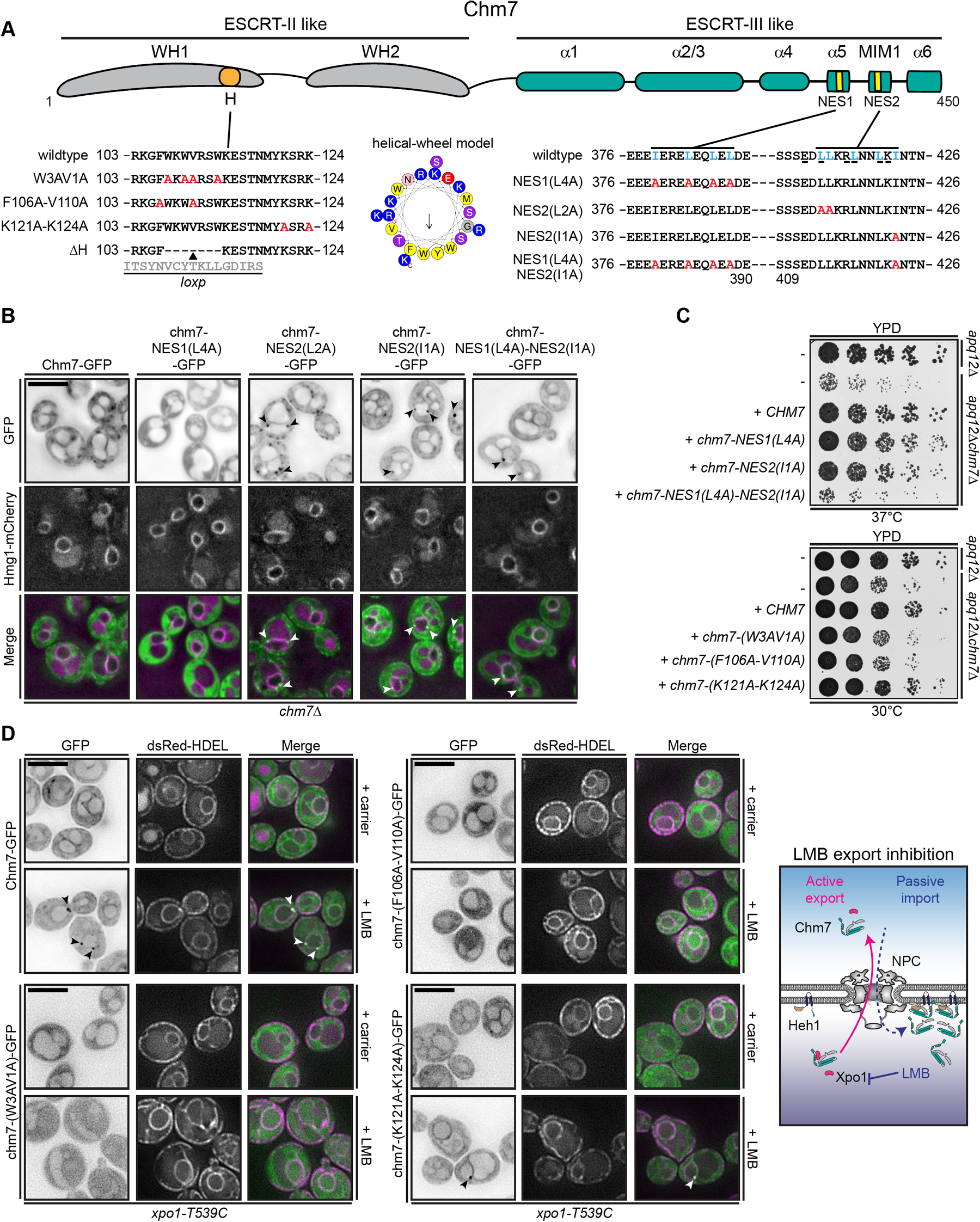
A conserved hydrophobic stretch and NES2 are required for Chm7 function. **(A)** Schematic of Chm7 with predicted winged helices (WH), hydrophobic stretch (H), alpha helices (α), MIT interacting motif type 1 (MIM1) and nuclear export signals (NES). Amino acid sequences encoded by alleles (with names at left) of *CHM7* with amino acid changes indicated in red in both the H and NES2/MIM1 sequences. In NES2/MIM1 sequence, hydrophobic residues that are predicted to be part of the NESs (Xu et al., 2015; Fung et al., 2017) are shown in blue whereas underlined residues are predicted to be important for MIM1 function (Obita et al., 2007; Stuchell-Brereton et al., 2007). In middle is a helical-wheel model of the hydrophobic stretch generated with HeliQuest (Gautier et al., 2008). Arrow indicates the hydrophobic moment. **(B)** NES2 is required to prevent accumulation of Chm7 at the INM. Deconvolved fluorescence micrographs of Chm7-GFP and indicated chm7-GFP proteins (fluorescence inverted) with Hmg1-mCherry as a nuclear envelope/ER marker, and merged images. Arrowheads point to GFP-foci at the nuclear envelope. Scale bar is 5 µm. **(C)** NES2 and the hydrophobic stretch are required for Chm7 function. Tenfold serial dilutions of the indicated strains with indicated *CHM7* alleles plated onto YPD and grown at indicated temperatures before imaging. **(D)** The hydrophobic stretch is required for the focal accumulation of Chm7 at the INM. Deconvolved fluorescence micrographs of Chm7-GFP and indicated chm7-GFP proteins (fluorescence inverted), HDEL-dsRed (demarking the nuclear envelope/ER) and merged images, in the LMB-sensitive strain (*xpo1-T539C*), after treatment with carrier (MeOH) or LMB. Scale bar is 5 µm. At right is a cartoon of interpretation of Chm7 localization upon LMB induced inhibition of nuclear export. Arrowheads point to GFP-foci at the nuclear envelope.

First, we focused on the two NESs. We abrogated NES1 by altering all four of its isoleucine and leucine residues to alanine (L4A) and evaluated the localization of a moderately overexpressed chm7-NES1(L4A)-GFP protein *in vivo*. Hmg1-mCherry was used to visualize the nuclear envelope. As shown in Fig. 1 B, chm7-NES1(L4A)-GFP localized throughout the cytosol with no obvious accumulation in the nucleus or at the nuclear envelope. Thus, NES1 did not appreciably alter Chm7 distribution, although we note that fewer cytosolic structures were visible compared to wildtype Chm7-GFP.

We next took a more conservative approach with NES2 as mutation of all hydrophobic residues in this region would also be predicted to abrogate the function of its putative MIM1 domain (Fig. 1 A; underlined residues; also see (Bauer et al., 2015; Webster et al., 2016; Thaller et al., 2019)). Consistent with this idea, mutation of the first two leucines in NES2 to alanine resulted in the focal accumulation of chm7-NES2(L2A)-GFP throughout the cell (Fig. 1 B). This phenotype was reminiscent of the distribution of Chm7-GFP that we had previously observed in *vps4Δ* backgrounds (Webster et al., 2016; Thaller et al., 2019) and is consistent with a disruption in MIM function. In marked contrast, mutation of the single isoleucine (NES2(I1A)), which is outside of the MIM1 domain but would nonetheless be predicted to impact Xpo1 binding (Fung et al., 2017), led to the specific accumulation of chm7-NES2(I1A)-GFP in foci at the nuclear envelope (Fig. 1 B). These data are thus consistent with the interpretation that NES2 is the functional NES and is required to restrict access of Chm7 to the nucleus and the INM. These data are also consistent with work in preprint showing the discovery of an NES in the human protein (Vietri et al., 2019), though the predicted MIM function of this domain may not be conserved (Gatta et al., 2020).

It is also worth noting that the combination of NES1(L4A) and NES2(I1A) mutations resulted in the accumulation of chm7-NES1(L4A)-NES2(I1A)-GFP on the nuclear envelope as expected, but it was qualitatively less focal and appeared more evenly distributed along the nuclear envelope when compared with chm7-NES2(I1A)-GFP (Fig. 1 B). Thus, taken together, we favor a model in which NES2 is the functional NES but that the NES1 sequence impacts the ability of Chm7 to focally-accumulate on membranes, perhaps by contributing to Chm7 activation. Consistent with this idea, this sequence aligns with the “linker” region just upstream of alpha helix 5 in the ESCRT-III, Snf7 (Webster et al., 2016), and which contributes to Snf7 activation (Henne et al., 2012). As such, both sequences contribute to Chm7 function as expression of neither chm7-NES2(L2A) nor chm7-NES1(L4A) could fully complement the synthetic growth defects of c*hm7Δapq12Δ* strains (compare colony sizes to *CHM7*-complementation; (Fig. 1 C)), and in fact an allele expressing both the NES1(L4A) and NES2(I1A) mutations was not functional in this context (Fig. 1 C).

We next tested how mutation of the hydrophobic stretch in the N-terminus of Chm7 impacted its distribution and function (Fig. 1 A). This region, which is not that well conserved at the sequence level (Fig. S1, A and B) nonetheless retains a hydrophobic character like its human counterpart, which imparts binding to ER membranes (Olmos et al., 2016). Interestingly, the region can be modeled as an alpha-helix: The orientation of the hydrophobic and charged residues is evocative of an amphipathic helix (Fig. 1 A and Fig. S1 B). We therefore generated mutations in the coding sequence of the hydrophobic stretch where codons for either hydrophobic (*chm7-(W3AV1A)* and *chm7-(F106A-V110A)*) or charged (*chm7-(K121A-K124A)*) amino acid residues were mutated to encode alanine. Previous work with the human protein identified residues essential for ER association within this same region (Olmos et al., 2016). However while overexpressed human CHMP7 localizes to the ER, yeast Chm7 does not bind to ER unless Heh1 localization to the INM is inhibited (Thaller et al., 2019). Therefore, in order to test whether these mutations affected Chm7’s membrane-association, we had to provide them access to the nucleus and to Heh1 at the INM.

To allow the Chm7 mutants access to the nuclear interior, we assessed the localization of Chm7-GFP, chm7-(W3AV1A)-GFP, chm7-(F106A-V110A)-GFP and chm7-(K121A-K124A)-GFP after inhibition of Xpo1 with Leptomycin B (LMB) in the LMB-sensitive *xpo1-T539C* strain (Neville and Rosbash, 1999) co-expressing a dsRED-HDEL fluorescent ER lumenal marker (Madrid et al., 2006). Consistent with our prior work (Thaller et al., 2019), LMB treatment results in the recruitment of Chm7-GFP into focal accumulations at the INM (Fig. 1 D and see diagram for interpretation). In striking contrast, chm7-(W3AV1A)-GFP and chm7-(F106A-V110A)-GFP clearly accessed the nucleoplasm but failed to accumulate at the INM (Fig. 1 D). This effect was specific for the hydrophobic mutations as chm7-(K121A-K124A)-GFP was able to accumulate at the INM similarly to the wildtype protein. Thus, the hydrophobicity of amino acids within this hydrophobic stretch is required for Chm7’s association with the INM. Consistent with the conclusion that these membrane interactions are critical for Chm7 function, only *chm7-(K121A-K124A)-GFP* was able to complement the fitness loss of *chm7Δapq12Δ* strains, whereas *chm7-(W3AV1A)-GFP* and *chm7-(F106A-V110A)-GFP* could not (Fig. 1 C).

### The hydrophobic stretch confers binding to the INM independently of Heh1

Although mutation of the hydrophobic residues within the N-terminus of Chm7 impacted its ability to accumulate at the INM, we could not discern whether these mutations simply broke an interaction with Heh1, or whether they reflected a distinct Heh1-independent INM-binding mechanism. To evaluate this, we examined the localization of chm7-N-GFP (Fig. 2 A), which lacks the ESCRT-III domain and NESs and can thus access the nucleus and INM. Interestingly, upon overexpression, two distinct localizations of chm7-N-GFP were obvious at the nuclear envelope (visualized with Nup170-mCherry): a discrete focal accumulation (arrowheads, Fig. 2 B) concomitant with a general even distribution of nuclear envelope fluorescence. We therefore tested whether these two distributions were dependent on Heh1 or its paralogue Heh2. Consistent with prior data (Webster et al., 2016), deletion of *HEH1* led to a complete loss of the focal accumulation of chm7-N-GFP, but surprisingly it had no discernable impact on the more diffuse nuclear rim fluorescence. This result was specific for *HEH1* as deletion of *HEH2* had no impact on either distribution; in fact if anything there was higher chm7-N-GFP fluorescence within foci of *heh2Δ* cells. Deletion of both *HEH1* and *HEH2* mirrored results of deletion of *HEH1* alone. Thus, even in the absence of these two established Chm7 binding partners at the INM (Webster et al., 2016; Thaller et al., 2019), there remained a specific membrane-binding activity associated with the N-terminus of Chm7. We therefore tested whether deletion of the hydrophobic stretch (chm7-N-ΔH-GFP; Fig. 2 A) could confer this additional function. Indeed, chm7-N-ΔH-GFP was distributed throughout the cytoplasm and nucleus with no detectable accumulation on any structure (Fig. 2 B). Thus, we conclude that the hydrophobic stretch contributes to a specific INM-binding activity of Chm7 that is independent of the LEM proteins.

**Figure 2.**
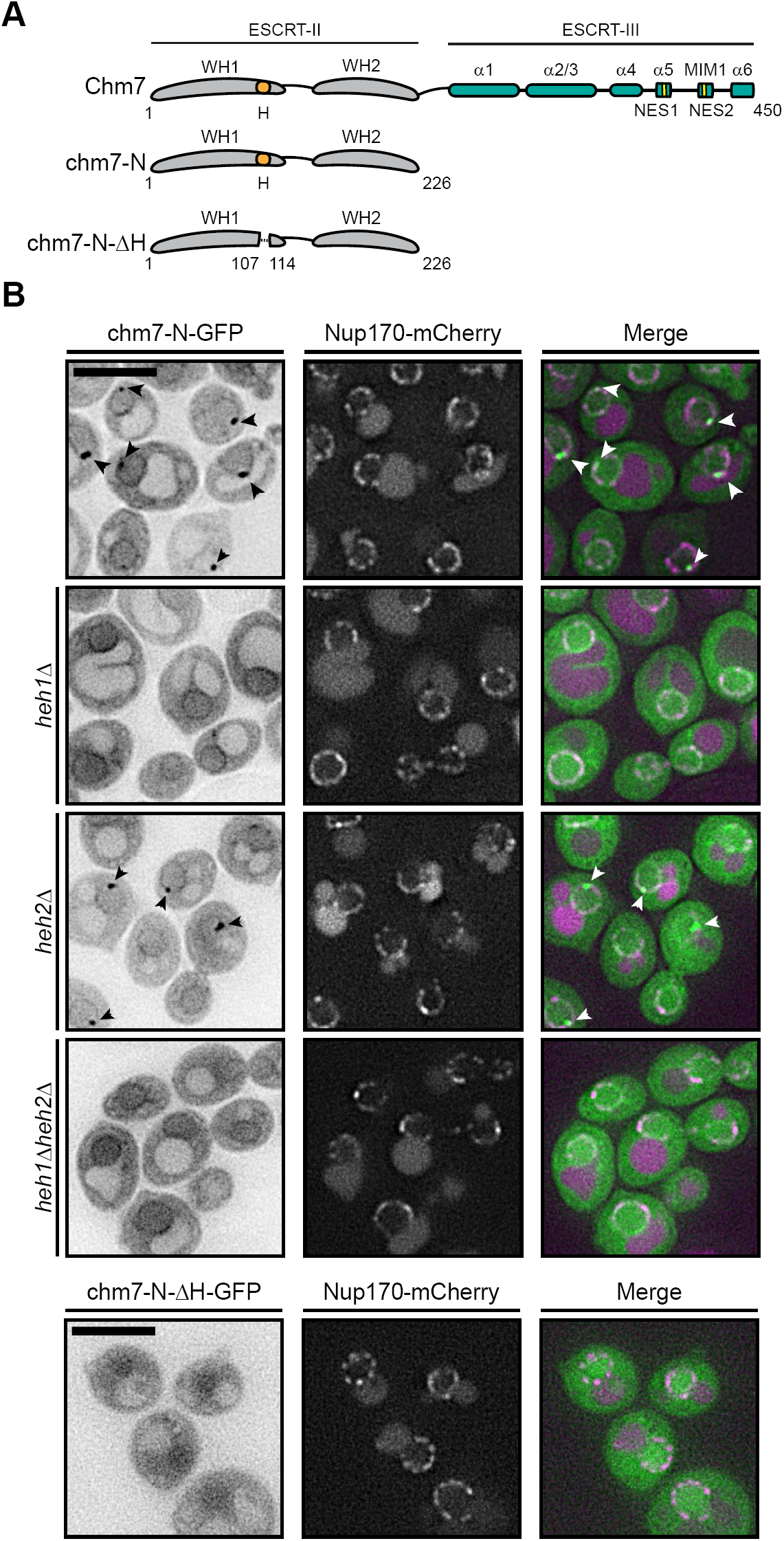
Chm7’s hydrophobic stretch confers Heh1-independent INM binding. **(A)** Schematic of Chm7 and indicated truncations. Numbers are amino acids. **(B)** Deconvolved fluorescence micrographs of indicated chm7-N-GFP constructs (inverted fluorescence) with Nup170-mCherry and merged images in the indicated strains. Arrowheads point to chm7-N-GFP foci at the nuclear envelope. Scale bars are 5 μm.

### Chm7 binds directly to PA-rich lipid bilayers

What was most curious about the localization of chm7-N-GFP was that even in the absence of Heh1, there remained a specific association with the nuclear envelope (Fig. 2 B). We surmised that this association was likely with the INM as, were it the ONM, one would expect a broader distribution throughout the cortical ER as well. What is the nature of this INM specificity, if not Heh1? Because of the importance of the hydrophobic residues within the hydrophobic stretch and its potential to form an amphipathic helix, we wondered whether it might in fact directly bind to specific lipids. To test this possibility, we generated recombinant GST-Chm7 and evaluated its ability to bind to a lipid strip containing a series of distinct phospholipid species (Fig. 3 A). Remarkably, compared to GST-alone, GST-Chm7 detected several anionic lipid species including phosphoinositides and phosphatidyl serine. Most noticeably, GST-Chm7 bound to PA and to cardiolipin. With the many established genetic relationships between PA-metabolism and the nuclear envelope (Santos-Rosa et al., 2005; Golden et al., 2009; Gorjánácz and Mattaj, 2009; Witkin et al., 2012; Bahmanyar et al., 2014; Grillet et al., 2016; Romanauska and Köhler, 2018; Barbosa et al., 2019; Penfield et al., 2020) and as cardiolipin is only found in mitochondria, we were particularly drawn to the potential that Chm7 might directly bind to PA.

**Figure 3.**
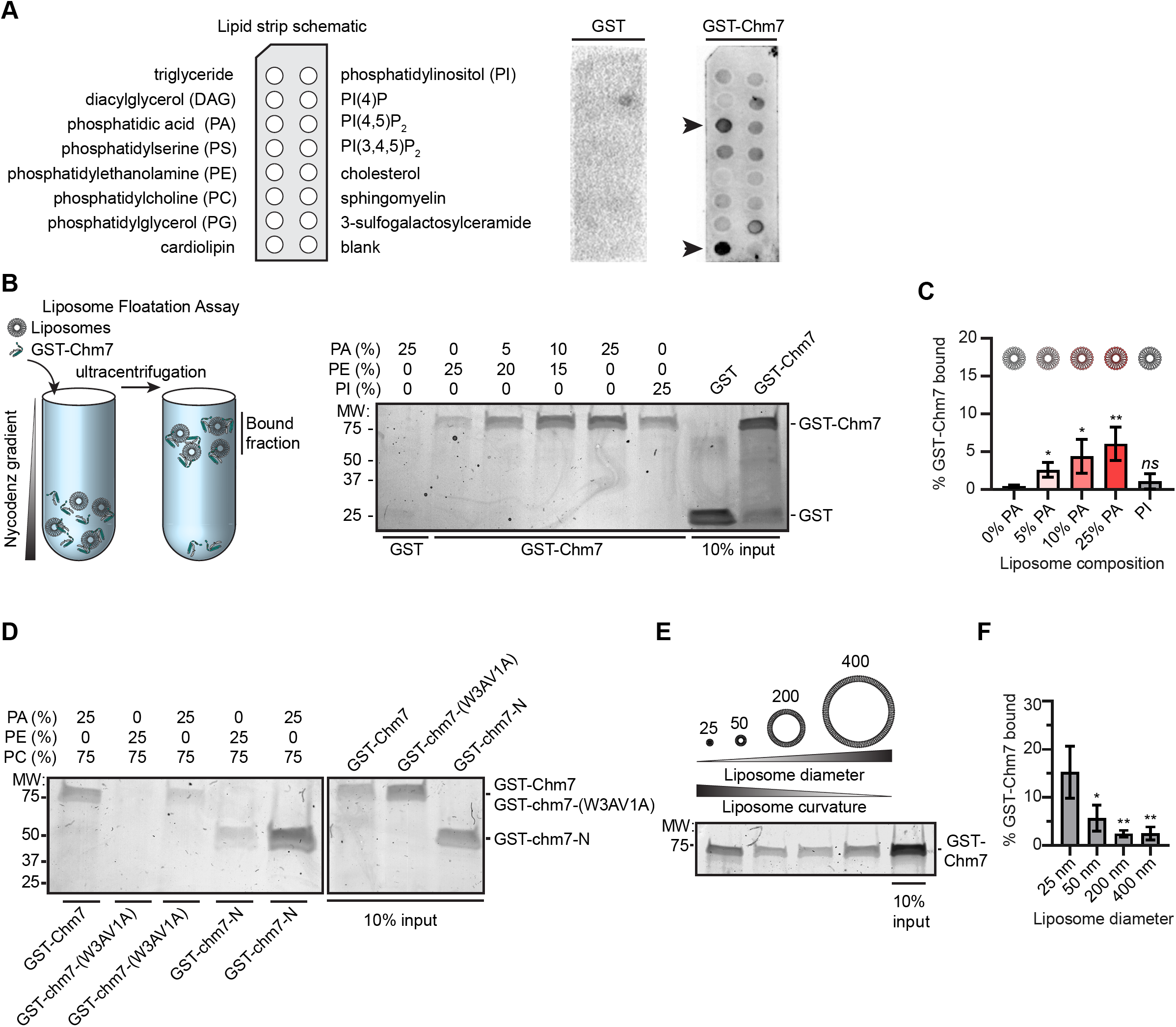
Chm7 binds directly to PA-rich lipid membranes. **(A)** GST-Chm7 preferentially binds to phosphatidic acid (PA) and cardiolipin. At left is schematic showing location of lipids immobilized on membranes (lipid strip). These lipid strips were probed with either GST (middle) or GST-Chm7 (right). Specific binding detected with anti-GST primary antibodies followed by HRP-conjugated secondary antibodies and ECL. PA and cardiolipin indicated with arrowheads. **(B)** GST-Chm7 specifically binds PA-rich liposomes. At left is schematic of liposome floatation assay in which proteins and liposomes of defined lipid compositions are mixed, overlaid with a Nycodenz gradient and then subjected to ultracentrifugation. Fractions that float (bound) were separated by SDS-PAGE and proteins detected with Coomassie staining; all liposomes are composed of 75% PC with the indicated percentages of PA, PE and PI. Numbers indicate the molecular weight (MW) in kD. **(C)** Histogram of quantification of the amount (relative to input) of GST-Chm7 bound to liposomes with increasing PA concentration. Error bars are SD from three independent experiments. *p* values were calculated from a one-way ANOVA with Tukey’s post-hoc test where *ns* is *p* > 0.05, * *p* ≤ 0.05, ** *p* ≤ 0.01. **(D)** Mutations in the hydrophobic stretch of Chm7 diminishes PA-binding. Coomassie-stained gel of liposome floatation and analysis as in (B) but incorporating GST-chm7-N and GST-chm7-(W3AV1A). **(E)** Chm7 prefers high curvature. Coomassie-stained gel of liposome floatation and analysis as in (B) but changing liposome diameter as indicated above liposome diagrams (in nm). All liposomes in this experiment are composed of 5% PA, 20% PE and 25% PC. **(F)** Histogram of quantification of the liposome-bound fraction of GST-Chm7 in (E) normalized to input. Error bars are SD from three independent experiments. *p* values were calculated from a one-way ANOVA with Tukey’s post-hoc test where * *p* ≤ 0.05, ** *p* ≤ 0.01.

As lipid strips do not mimic the physiological environment of a lipid bilayer, we next incubated recombinant GST-Chm7 (or GST alone) with liposomes generated with increasing concentrations of PA, from 5 to 25%. As diagramed in Fig. 3 B, these liposome-protein mixtures were then overlaid with a Nycodenz gradient and subjected to ultracentrifugation. Under these conditions, liposomes with bound proteins float; floated fractions are then separated by SDS-PAGE and visualized with coommassie staining. Gratifyingly, as we increased the amount of PA, we observed a commensurate increase in GST-Chm7 bound to the PA-rich liposomes supporting that Chm7 binds directly to PA (Fig. 3, B and C). This conclusion is further supported by the observation that Chm7 shows a weaker preference for PI-rich liposomes (which share a similar charge as PA) and for PE-rich liposomes (which share a similar cone-shape)(Van Meer et al., 2008). Consistent with the idea that this binding is mediated by the hydrophobic stretch, GST-chm7-N also shows, perhaps more efficient, binding to PA-rich liposomes and mutation of hydrophobic residues (W3AV1A) within the hydrophobic stretch in the context of the full length protein, abrogates (but does not fully abolish) this interaction (Fig. 3 D). Thus, we conclude that Chm7 has a preference for binding to PA, and this interaction requires the hydrophobic stretch in its N-terminus.

We next also assessed whether Chm7 might have a preference for binding to flat or highly curved membranes. To test this, we generated liposomes with different diameters (from ∼400 nm to ∼25 nm) and tested their ability to float GST-Chm7. As shown in Fig. 3 E and quantified in Fig. 3 F, GST-Chm7 had a clear preference for both ∼50 nm and ∼25 nm diameter liposomes compared with those that were essentially flat (i.e. ∼200 and ∼400 nm). Thus, taken together, both lipid composition and curvature contribute to the efficiency by which Chm7 is recruited to membranes *in vitro*.

### Global changes to PA-levels disrupt nuclear-cytoplasmic compartmentalization

Considering that Chm7 bound to PA *in vitro*, we wondered how altering global PA-levels in cells would impact Chm7 distribution *in vivo*. We therefore tested the localization of endogenously-expressed Chm7-GFP in strains where the gene encoding the lipin orthologue, *PAH1*, was deleted; *pah1Δ* cells show a two-fold increase in total PA levels (Han et al., 2006), including an increase at the INM (Romanauska and Köhler, 2018), alongside changes to nuclear morphology (Santos-Rosa et al., 2005). As shown in Fig. 4 A, in *pah1Δ* strains, endogenously-expressed Chm7-GFP redistributed from the cytosol to membranes throughout the cell including the nuclear envelope (Fig. 4 A, arrowheads) and vacuole membrane (Fig. 4 A, asterisks). Thus, increased levels of PA impacts Chm7’s distribution, further re-enforcing that elevating bilayer PA is an input to its recruitment to a membrane interface. Interestingly, Chm7’s recruitment to the nuclear envelope may also reflect an activation of nuclear envelope surveillance as *CHM7* was required for *pah1Δ* viability (Fig. 4 B, and Fig S2 A). Indeed, this fitness loss was likely a reflection of disturbing nuclear-cytoplasmic compartmentalization as we observed that *pah1Δ* cells, particularly *chm7Δpah1Δ* cells, were unable to effectively accumulate an NLS-GFP reporter in a subset of cells (Fig. 4 C, and Fig. S2 B). Therefore, Chm7 is protective of nuclear-cytoplasmic compartmentalization in the context of elevated PA, which is itself detrimental to membrane integrity (Kooijman et al., 2003; Kwolek et al., 2015). These data fit well with recent reports that changes to lipid metabolism might negatively impact nuclear envelope integrity including lipid droplet biogenesis at the INM (Barbosa et al., 2019).

**Figure 4.**
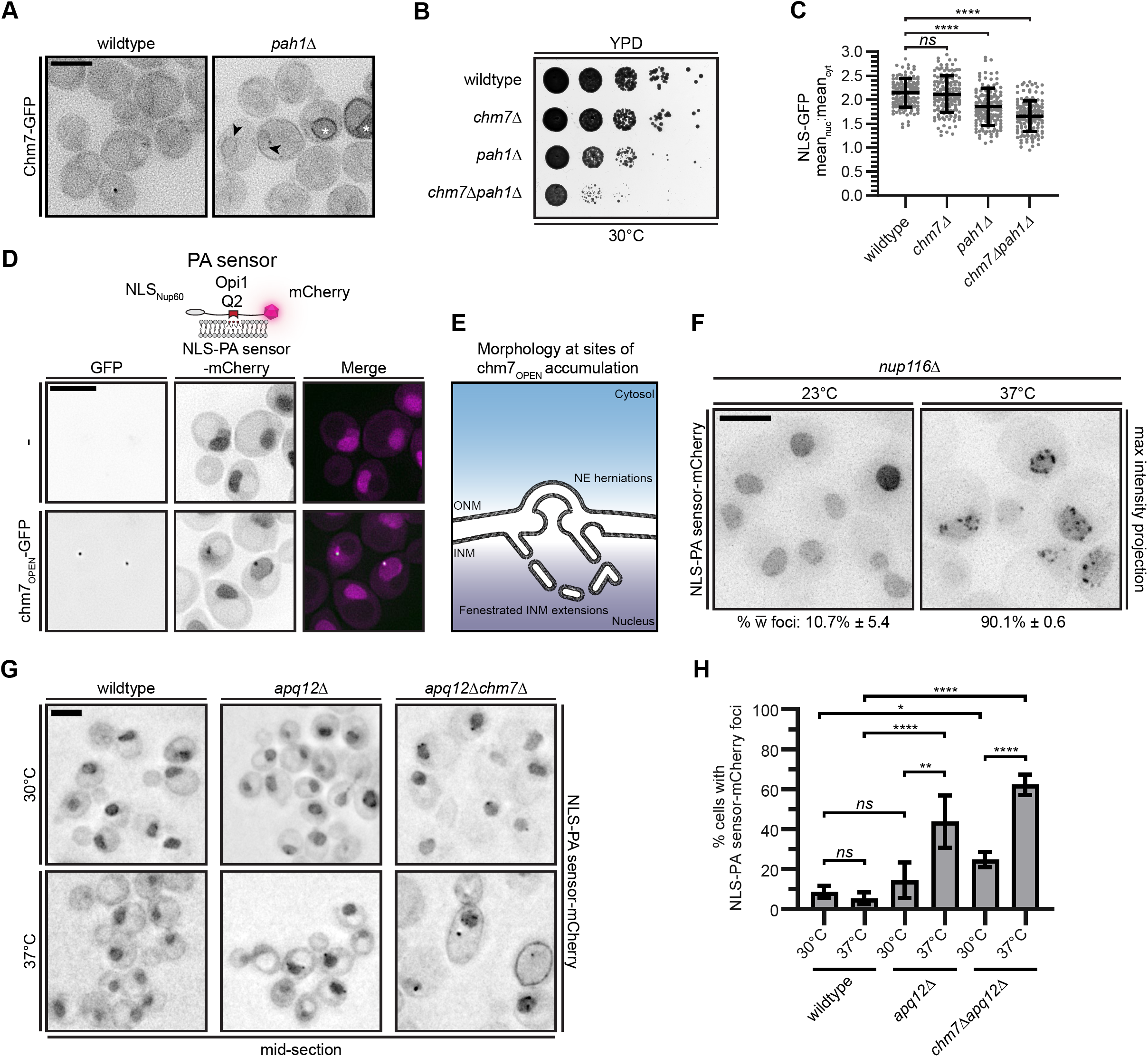
Recruitment of Chm7 to the nuclear envelope drives local PA accumulation. **(A)** Changes to global PA-levels leads to Chm7 recruitment to membranes. Deconvolved inverted fluorescence micrographs of Chm7-GFP in wildtype and *pah1Δ* strains. Arrowheads indicate Chm7-GFP accumulation at the nuclear envelope and asterisks denote Chm7-GFP vacuole-membrane localization. Scale bar is 5 µm. **(B)** *CHM7* is required to maintain fitness of *pah1Δ* cells. Tenfold serial dilutions of the indicated strains grown at 30°C for 48 h prior to imaging. **(C)** Nuclear-cytosolic compartmentalization is perturbed in *pah1Δ* cells. Scatter plot of the mean nuclear (nuc) to cytosolic (cyt) fluorescence intensity with SD from three independent experiments (50 cells/strain/experiment) of an NLS-GFP reporter from cells of the indicated genotype cultured at 23°C. Images from a representative experiment shown in Fig. S2 B. *p* values were calculated from a one-way ANOVA with Tukey’s post-hoc test where *ns* is *p* > 0.05, and **** *p* ≤ 0.0001. **(D)** PA accumulates at site of Chm7 hyperactivation. Deconvolved fluorescence micrographs of cells with no GFP (-) or chm7_OPEN_-GFP with NLS-PA sensor-mCherry, which binds to PA (see diagram at top). GFP (inverted), mCherry (inverted), and merged images are shown. **(E)** Cartoon of local nuclear envelope morphology at sites of chm7_OPEN_ accumulation (described in Thaller et al., 2019) including nuclear envelope herniation and the expansion of a fenestrated network of membranes that emanate from the INM. **(F)** PA focally accumulates at the nuclear envelope in strains with NPC assembly-associated herniations. Deconvolved inverted fluorescence images (maximum intensity projections of a z-series) of NLS-PA sensor-mCherry in a *nup116Δ* strain at 23°C (left) or after 3 h at 37°C, which triggers herniation formation (right). Numbers are average percentage and SD of cells from three independent experiments with the appearance of NLS-PA sensor-mCherry foci at the nuclear envelope. **(G)** Deconvolved fluorescence micrographs of NLS-PA sensor-mCherry in the indicated strains after 3 h at 30°C (upper panels) or 37°C (lower panels). The average percentage of cells and SD from three replicates with the appearance of NLS-PA sensor-mCherry foci at the nuclear envelope and are plotted in (H). **(H)** Histogram of quantification of the mean and SD of cells with the appearance of NLS-PA sensor-mCherry foci at the nuclear envelope from three independent experiments analogous to (G). > 50 cells were quantified per experiment. *p* values were calculated from a one-way ANOVA with Tukey’s post-hoc test where *ns* is *p* > 0.05, * *p* ≤ 0.05, ** *p* ≤ 0.01, **** *p* ≤ 0.0001.

### Chm7 hyperactivation drives local PA accumulation at the INM

To further explore the functional and physical relationship between Chm7 and PA, we tested whether we could directly observe PA accumulation at sites of Chm7 activation. We therefore took advantage of a recently developed PA sensor that was generated from the PA-binding domain of Opi1 (Romanauska and Köhler, 2018). To test PA levels at the INM, this sensor is fused to an NLS and mCherry to visualize its distribution. As there is very little PA at the INM under normal conditions, this PA sensor localizes throughout the nucleoplasm (Fig. 4 D; also see (Romanauska and Köhler, 2018)). In contrast, a DAG-sensor generated with the PKCβ DAG binding domain with an NLS (Romanauska and Köhler, 2018) localizes to the INM and can also be visualized at other membranes throughout the cell (Fig. S2 C; (Romanauska and Köhler, 2018)).

With these tools in hand, we first tested whether expression of a hyperactivated form of Chm7, chm7_OPEN_, which is constitutively localized into one or two foci at the INM (Webster et al., 2016; Thaller et al., 2019), impacts either sensor’s distribution. A reminder that in prior work, we showed that at sites of chm7_OPEN_ accumulation, there is an expansion of a fenestrated cisternal network of membrane emanating from INM, in addition to nuclear envelope herniations (diagrammed in Fig. 4 E; also see (Thaller et al., 2019)). Remarkably, upon expression of chm7_OPEN_, we observed the appearance of the PA sensor, but not the DAG sensor, in a focus at the periphery of the nucleus (Fig. 4 D, and Fig. S2 C). As this focus perfectly colocalized with chm7_OPEN_-GFP, we interpret these data in a model in which chm7_OPEN_ directly recruits PA to this domain at the INM, or there are local changes to PA metabolism at sites of chm7_OPEN_ accumulation.

To begin to differentiate between these possibilities, we investigated whether we could detect analogous changes to PA-distribution at the INM under more physiological conditions. For example, we had previously shown that Chm7 accumulates at the nuclear envelope upon perturbation of *de novo* NPC assembly (Webster et al., 2016). This recruitment is well correlated with the presence of nuclear envelope herniations associated with malformed NPCs observed in both *apq12Δ* (Scarcelli et al., 2007) and *nup116Δ* (Wente and Blobel, 1993) genetic backgrounds; in fact, both of these genetic backgrounds require *CHM7* for viability (Bauer et al., 2015; Webster et al., 2016). These strains also have the additional advantage that they are temperature sensitive and form herniations upon shift from 23°C or 30°C to higher (37°C) temperatures (Wente and Blobel, 1993; Scarcelli et al., 2007; Webster et al., 2016).

We first examined PA sensor distribution in *nup116Δ* cells grown at (23°C) or at herniation-forming (37°C) temperatures. Importantly, at 23°C, the PA sensor localized in the nucleus as expected with no detectable accumulation at the INM (Fig. 4 F). Upon shifting the culture to 37°C, there was a striking re-distribution of the PA sensor, which now accumulated along the nuclear periphery in distinct foci in 90% of cells (Fig. 4 F). Importantly, this PA sensor accumulation was not due to the temperature shift, per se, as wildtype cells did not exhibit this phenotype (Fig. 4 G). Similar results were observed in *apq12Δ* cells, although it was less penetrant with a maximum of ∼40% of cells showing a focal accumulation at the nuclear periphery (Fig. 4, G and H). To our knowledge, this is the first demonstration of such a local accumulation of PA at the INM and strongly suggests that PA is either being generated at these sites and/or its diffusion along the lipid bilayer is restricted. Importantly, recruitment of the PA sensor to membranes is insufficient to drive local PA accumulation (Romanauska and Köhler, 2018). Thus, with this in mind, we favor an interpretation in which PA is being locally enriched at the INM due to an intrinsic change in the properties of the membrane itself, which is logically imparted by the herniations.

As a dominant form of Chm7 was sufficient to drive the focal accumulation of PA (Fig. 4 D), it was a reasonable hypothesis that Chm7 recruitment to the nuclear envelope, which we previously observed in *nup116Δ* and *apq12Δ* cells (Webster et al., 2016), could contribute to this focal PA sensor distribution. We therefore tested whether the PA accumulation observed in *apq12Δ* strains was a cause or consequence of Chm7 recruitment by monitoring PA sensor distribution in *apq12Δchm7Δ* strains. As shown in Fig. 4 G, the absence of Chm7 did not appreciably impact the appearance of nuclear envelope-associated foci, in fact, a higher percentage (60%) of cells had visible foci (Fig. 4 H). Thus, PA sensor focal-accumulation does not require Chm7, suggesting that local PA-accumulation at the INM is likely upstream of Chm7 recruitment.

Of further interest, we also observed a dramatic change in PA sensor distribution in a population of *apq12Δchm7Δ* cells such that it localized to the cell cortex (Fig. 4 G). We think that it is most likely that this phenotype is a result of nuclear rupture, as we have observed a loss of nuclear-cytoplasmic compartmentalization of an NLS-GFP reporter in *apq12Δchm7Δ* cells (Webster et al., 2016) coincident with discontinuities in the nuclear envelope at the level of ultrastructure (Thaller et al., 2019). We further note that in these cortical membranes, the PA sensor is no longer focal suggesting that there is a loss of the underlying membrane structure that leads to PA accumulation under these conditions.

### PA sensor recruitment is likely at nuclear envelope herniations

Cumulatively then, we conclude that the changes in local PA distribution in *nup116Δ* and *apq12Δ* cells are not caused by Chm7, but instead by the upstream insult that leads to Chm7 recruitment. We therefore hypothesized that PA was in fact accumulating at sites of NPC misassembly, perhaps within the herniations themselves. To explore this possibility, we first tested whether we could observe a spatial relationship between the PA sensor and NPCs and/or the likely defective NPC assembly intermediates at the bases of the herniations (Wente and Blobel, 1993; Thaller et al., 2019; Allegretti et al., 2020). We therefore localized the PA sensor in *nup116Δ* strains that co-expressed GFP-Nup49. As expected, we observed minimal coincidence between the localization of the PA sensor and GFP-Nup49 at the permissive temperature (Fig. 5 A, top panels). This is evident in line profiles of both the mCherry and GFP fluorescence along the nuclear envelope of a single cell (boxed cell, Fig. 5 A) and in a scatterplot showing no correlation (*r* = 0.1143) between the PA sensor and GFP-Nup49 signals along nuclear envelopes derived from multiple cells (Fig. 5 A, top, far right). However, under conditions in which herniations form (37°C), we observed a clear co-localization of the mCherry and GFP signals as shown in the overlap of the GFP and mCherry peaks in the line profile shown (Fig. 5 A, bottom panels). Further, there is an obvious increase in the correlation of the mCherry and GFP signals with an *r* = 0.6220 (Fig. 5 A, bottom, far right). This result was then consistent with our hypothesis and suggested that PA may in fact accumulate at or near sites of NPC biogenesis-associated nuclear envelope herniations.

**Figure 5.**
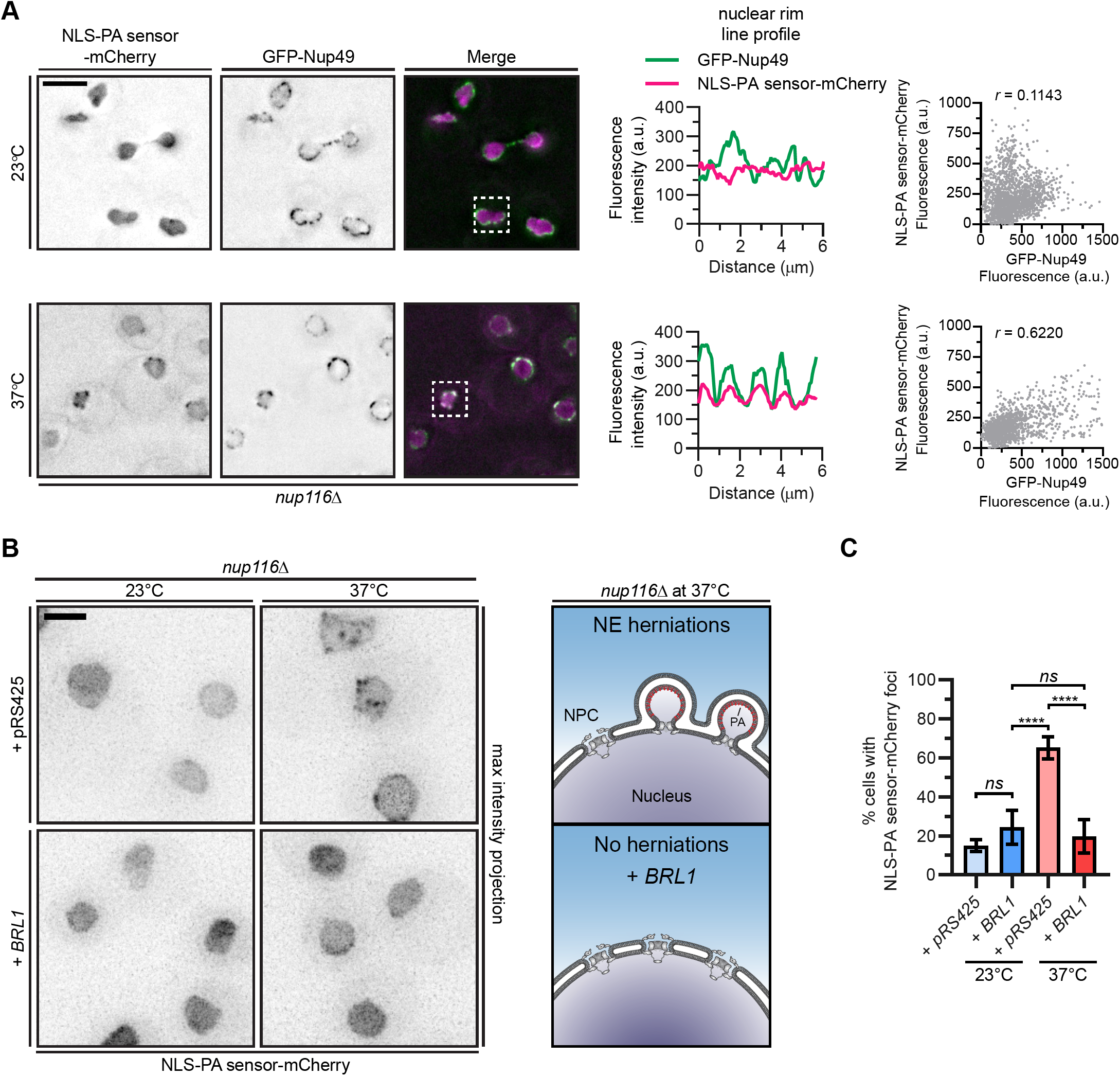
PA accumulates at nuclear envelope herniations associated with NPC assembly defects. **(A)** The PA sensor co-localizes with nups under conditions of herniation formation. Deconvolved fluorescence micrographs of *nup116Δ* cells expressing NLS-PA sensor-mCherry and GFP-Nup49 at the indicated temperatures. In the GFP and mCherry images the fluorescence is inverted whereas the merge shows the GFP signal in green and mCherry in magenta. Scale bar is 5 µm. Line profiles of the fluorescence intensities (in arbitrary units, a.u.) along the nuclear rim of the boxed cells are shown at right of images. At far right, correlation of fluorescence intensity (in a.u.) of GFP-Nup49 and NLS-PA sensor-mCherry from line profiles drawn around the nuclear envelope. Linear regression calculated from 15 randomly selected cells; *r* is the linear correlation coefficient (Pearson’s). **(B)** The PA sensor fails to accumulate at the nuclear envelope when herniations are genetically resolved. Maximum intensity projections of a deconvolved z-series of inverted fluorescence images of NLS-PA sensor-mCherry in a *nup116Δ* strain transformed with either pRS425 or pGP564, which contains a genomic fragment including *BRL1, PIH1, YHR035W, PUT2, RRF1, MSC7, VMA10, BCD1*, and *SRB2*. Cells were cultured at either at 23°C or 37°C. Interpretation in cartoon at right. Scale bar is 2.5 µm. **(C)** Histogram with mean and SD showing the percentage of cells with NLS-PA sensor-mCherry nuclear rim foci from three independent experiments as in (C). Error bars plot the SD from three independent experiments of > 50 cells. *p* values were calculated from a one-way ANOVA with Tukey’s post-hoc test where *ns* is *p* > 0.05, **** *p* ≤ 0.0001.

### Disruption of nuclear envelope herniations prevents PA accumulation

Lastly, to provide additional support that the local accumulation of PA may in fact be due to the presence of nuclear envelope herniations, we took advantage of recent work that demonstrated that *nup116Δ* nuclear envelope herniations could be resolved by overexpressing *BRL1* (Zhang et al., 2018). *BRL1* is an essential integral nuclear envelope membrane protein that localizes to sites of NPC assembly and nuclear envelope herniations; it has been proposed to contribute to INM-ONM fusion (Zhang et al., 2018). As shown in Fig. 5 B and C, the overexpression of *BRL1* dramatically reduced the penetrance of the focal accumulations of the PA sensor (from 65 to 19%) at the herniation-forming temperature. Thus, under conditions in which there are fewer nuclear envelope herniations, there is a coincident reduction in PA sensor accumulation at the INM. When taken in the context of the localization of the PA sensor at NPCs, and the need for a mechanism to restrict PA-diffusion (logically provided by the negative curvature of the herniations themselves; (Domanov et al., 2011)), we are strongly in favor of a model in which PA accumulates within the INM of the herniation structure.

### Summary and outlook

A model in which PA accumulates at sites of NPC assembly-associated nuclear envelope herniations has many potential implications. First, it suggests that the accumulation of PA in herniations may contribute to herniation growth and be part of the pathology associated with these structures. Such an idea fits well with work exploring the mechanism behind the biogenesis of similar herniations observed in cells with a loss of function of the AAA+ ATPase, TorsinA (Goodchild et al., 2005; Laudermilch et al., 2016). As data continues to grow linking these herniations to defective NPC assembly (Laudermilch et al., 2016; Pappas et al., 2018; Rampello et al., 2020), so does evidence that they could be a product of defective lipid metabolism; in fact recent work suggests that TorsinA may inhibit Lipin (Grillet et al., 2016; Cascalho et al., 2019). Second, and as follows, while ultimately too much PA may be deleterious (Oliveira et al., 2010; Adeyo et al., 2011; Park et al., 2015), it seems probable that a tightly controlled system of PA generation may actually promote INM-ONM fusion during interphase NPC assembly. Such an idea would be consistent with the emerging role of PA in several other membrane fusion events, e.g. COPI vesicle formation (Yang et al., 2008; Park et al., 2019) and exocytosis (Zeniou-Meyer et al., 2007; Miner et al., 2017) among many others (Corrotte et al., 2006; Nakanishi et al., 2006; Liu et al., 2007; Yang et al., 2008; Giridharan et al., 2013; Bates et al., 2014; Adachi et al., 2016; Starr et al., 2016), and could ultimately underlie genetic relationships defined between PA-metabolism and nup genes (Siniossoglou et al., 1998; Tange et al., 2002; Santos-Rosa et al., 2005; Costanzo et al., 2016; Lord and Wente, 2020) and those that link lipid metabolism more generally to nuclear envelope function (Gorjánácz and Mattaj, 2009; Domart et al., 2012; Bahmanyar et al., 2014; Grillet et al., 2016; Romanauska and Köhler, 2018; Ventimiglia et al., 2018; Kinugasa et al., 2019; Barbosa et al., 2019; Kume et al., 2019; Lee et al., 2020; Penfield et al., 2020). Definitively testing such a model requires more effective tools to precisely toggle PA levels with nanometer precision; recently developed light-inducible tools might provide a way forward here in the future (Tei and Baskin, 2020).

This apparent dichotomy where too much PA may be deleterious and just the right amount productive likely also extends to the role of PA in nuclear envelope surveillance and repair. Indeed, as shown here, increased levels of PA driven by deletion of *PAH1* is itself a background where nuclear-cytoplasmic compartmentalization is lost (Fig. 4 C, and Fig. S2 B), likely due to nuclear envelope rupture, whereas PA-binding by Chm7 is essential for its function in the context of such ruptures (Fig. 1 C). The question remains: Why does Chm7 require PA-binding for its function? Here again, several non-mutually exclusive scenarios can be considered. The first is that whether a nuclear envelope rupture occurs during NPC biogenesis or at herniations (those associated with NPC assembly, and those that aren’t), a prediction would be the formation of a highly curved membrane where the INM-ONM would likely spontaneously come back together after rupture. PA would have a natural affinity for these destabilized membranes and may thus provide a reinforcing targeting signal for Chm7. This makes sense as well in the context of our observed high curvature affinity of Chm7 (Fig. 3 E). Secondly, as a putative Chm7-Heh1 polymer forms (predicated after likely similarities with the analogous CHMP7-LEM2 interaction; (von Appen et al., 2020)), it may be possible to directly recruit additional PA (as seen in Fig. 4 D); this may have the added benefit of promoting the ultimate annular fusion event necessary for nuclear envelope sealing. Similarly, membrane remodeling by ESCRTs may itself increase local membrane tension (Booth et al., 2019) and trigger a local lipid synthesis response, which may contribute membrane to help seal nuclear ruptures (Kinugasa et al., 2019) and the nuclear envelope at the end of mitosis (Lee et al., 2020; Penfield et al., 2020).

Our initial model of nuclear envelope surveillance posited that the exposure of nuclear contents, specifically Heh1, to cytosolic Chm7 triggered local Chm7 activation and ultimately repair (Thaller et al., 2019). PA-binding must now be incorporated into this model as without the PA-binding hydrophobic stretch, we no longer observe an Heh1-dependent recruitment of Chm7 to the INM (Fig. 2 B). Thus, PA acts upstream or alongside Heh1 to recruit and activate Chm7. In a model in which Chm7 ultimately responds to the exposure of the INM to the cytosol, the incorporation of PA into this surveillance mechanism may in fact be used to prevent aberrant activation in the ER where PA levels are low and where newly synthesized Heh1 must travel to reach the INM. It follows then that there may be an inherent asymmetry in PA levels at the INM versus the ONM, even at steady state. Indeed, such a postulate could explain why chm7-N has a preference for binding the INM over the rest of the ER (Fig. 2 B). Thus, a goal going forward will be to develop methods to evaluate lipid composition specifically at the INM and, in so doing, more clearly define its function in NPC biogenesis, nuclear envelope surveillance and other, yet-to-be defined mechanisms.

**Figure S1.**
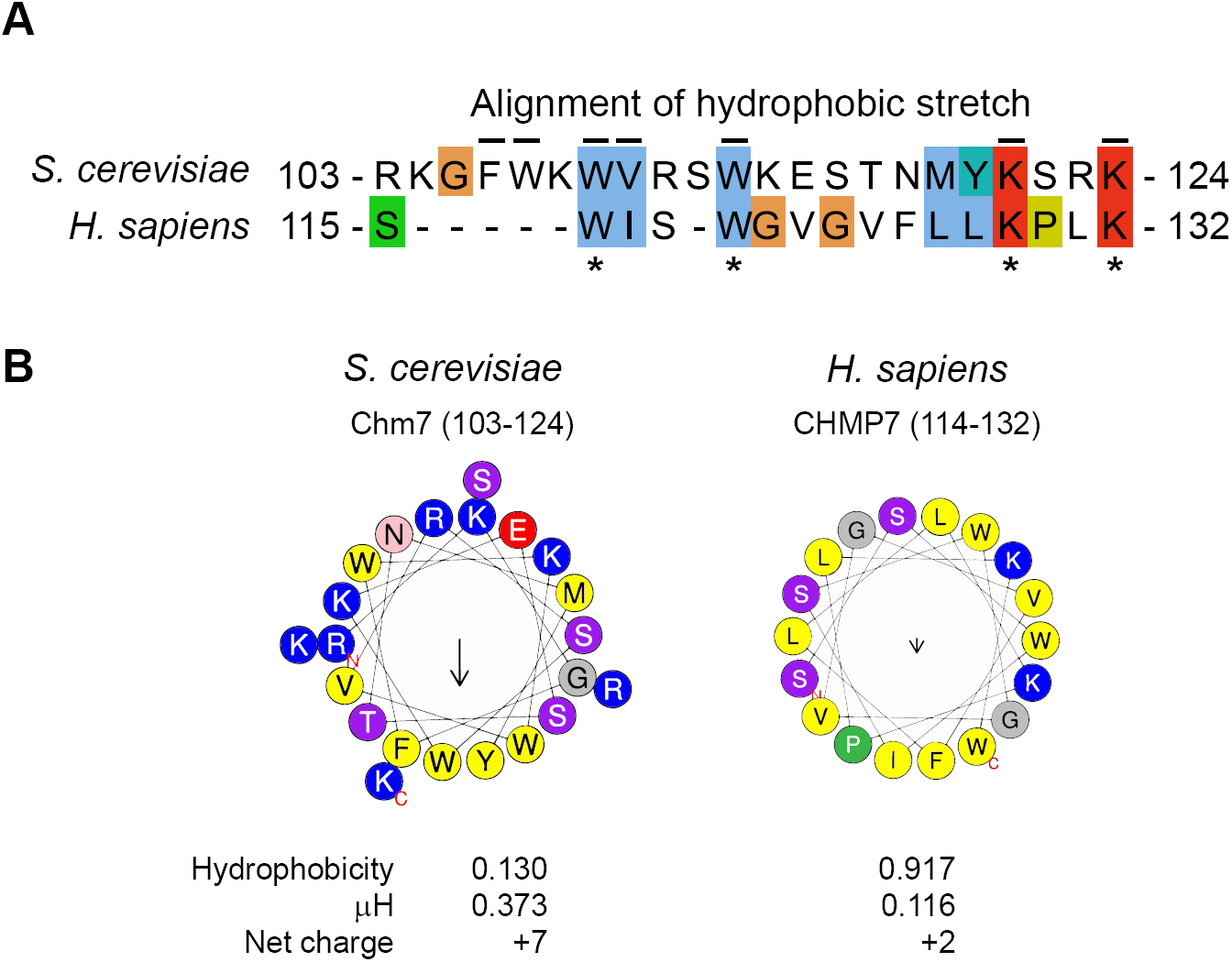
(supplement to Figure 1). Conserved hydrophobic stretch within Chm7 may be an amphipathic helix. **(A)** Alignment of hydrophobic stretches in budding yeast and Human. Color coding of properties of amino acids with conserved attributes as in ClustalW (Larkin et al., 2007) where hydrophobic residues are in blue, positive charged residues are red, negative charged residues are magenta, polar residues are green, glycines are orange, prolines are yellow, and aromatic residues are cyan. Numbers are amino acids. Asterisks denoted conserved residues, black lines are residues altered in mutational analysis in this study for Chm7 (top), and numbers are amino acids. **(B)** Helical-wheel diagrams, letters are amino acids. Physicochemical properties were calculated in HeliQuest (Gautier et al., 2008).

**Figure S2.**
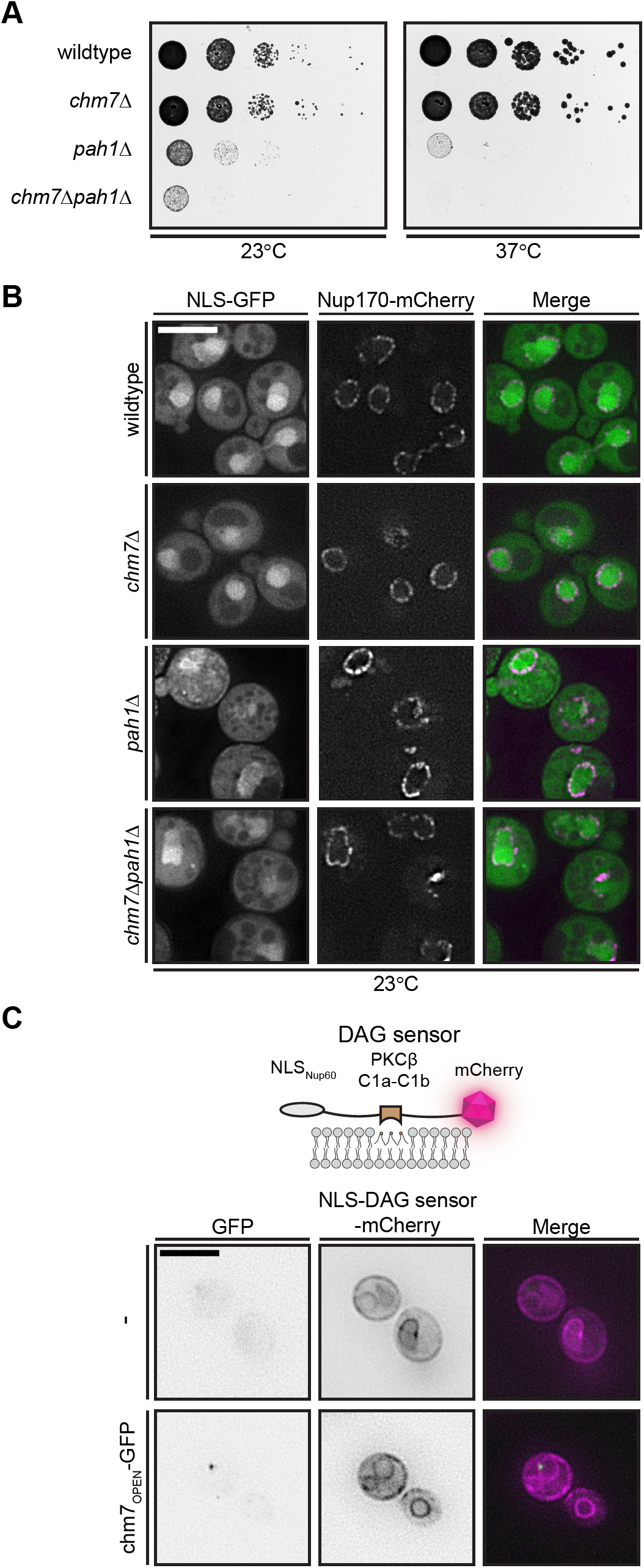
(supplement to Figures 4 and 5). Elevated membrane PA disrupts nuclear-cytoplasmic compartmentalization; Chm7 does not colocalize with a DAG sensor. **(A)** *CHM7* is required for fitness of *pah1Δ* strains. Tenfold serial dilutions of the indicated strains grown on YPD plates at 23°C or 37°C for 48 h before imaging. **(B)** Nuclear-cytoplasmic compartmentalization is perturbed in *pah1Δ* and c*hm7Δpah1Δ* cells. Deconvolved fluorescence micrographs of NLS-GFP, Nup170-mCherry and merge, in the indicated strains cultured at 23°C. **(C)** DAG does not accumulate at sites of Chm7 hyperactivation. Deconvolved fluorescence micrographs of cells with no GFP (-) or chm7_OPEN_-GFP with an NLS-DAG sensor-mCherry (See diagram at top). GFP (inverted), mCherry (inverted), and merge are shown. Scale bar is 5 µm.

---

Table S1. **List of genotype and origin of all *S. cerevisiae* strains used in this study**.

Table S2. **List of all plasmids used in this study**.

## Materials and Methods

### S. cerevisiae *culturing methods and strain generation*

For all experiments, cells were grown to mid-log phase in YPA (1% Bacto-yeast extract (BD), 2% Bacto-peptone (BD), 0.025% adenine hemi-sulfate (Sigma)) supplemented with 2% raffinose (R; BD), 2% D-galactose (G; Alfa Aesar) or 2% D-glucose (D; Sigma) or in Synthetic Complete (SC) medium lacking the indicated amino acids. The majority of experiments were performed at 30°C unless otherwise indicated. Standard protocols were followed for transformation, mating, sporulation, and dissection as described in (Amberg et al., 2005).

All yeast strains used in this study are listed in Table S1. Genetic modifications to generate gene deletions or the in-frame incorporation of fluorescent protein genes was accomplished using a PCR-based approach using the pFA6a plasmid series or pK3F (Table S2) as templates (Longtine et al., 1998; Zhang et al., 2017). Further, to generate DTCPL2030 loxP/Cre-mediated gene disruption was used to delete the coding sequence for the hydrophobic stretch (H) within the endogenous *CHM7* gene. Briefly, a kanamycin resistance cassette (*kan-MX6*) flanked by loxp sites was amplified off of pK3F (Zhang et al., 2017) using primers with 60 bp homology arms that flank the region targeted for deletion. The PCR product was transformed into DTCPL515 (in this strain the *CHM7* locus is modified to express *chm7-N-GFP* behind the *GAL1* promoter) and kanamycin-resistant colonies were selected. To generate the *GAL1-chm7-N-ΔH-GFP* allele, a colony containing the *KAN-MX6* cassette in the proper location was transformed with pSH47 (Güldener et al., 1996), which expresses the Cre recombinase under the control of the *GAL1* promoter. To induce Cre production, transformants were grown in YPAG for at least 4 h before plating to single colonies on YPD plates. Colonies were screened for loss of *KAN-MX6* and re-acquisition of fluorescence from *chm7-N-ΔH-GFP* after growth on YPAG.

To integrate NLS-PA sensor-mCherry into strains used for the temperature shift experiments, W303, *apq12Δ* (PCCPL249), *chm7Δapq12Δ* (CPL1323) and *nup116Δ* (Y1499) strains, pDT29 was linearized with *Stu*I (New England BioLabs) and genomically integrated at the *URA3* locus.

To generate strains to test complementation of *chm7Δapq12Δ*, a *chm7Δapq12Δ* (CPL1323) strain was transformed with *StuI* (New England BioLabs linearized pDT30, pDT31, pDT32, pDT33, pDT40, pDT41, or pDT44. Linearized plasmids were integrated at the *URA3* locus. Individual transformants were picked and GFP fluorescence was confirmed by microscopy after incubation overnight in YPAD.

### Plasmid generation

All plasmids used in this study are listed in Table S2.

To generate pDT30, an ORF encoding *CHM7-3xHA-GFP* was PCR amplified from genomic DNA isolated from DTCPL1240 and assembled into pRS406-GAL1 digested with *Eco*RI*/Xba*I (New England BioLabs) using the Gibson Assembly Master Mix (New England BioLabs).

Site-directed mutagenesis of NES and hydrophobic-stretch coding sequences was performed using the Q5 site-directed mutagenesis kit (New England Biolabs) with either pCPLJJ50, pDT30 or pDT40 as templates. Mutations were confirmed by sequencing.

pDT29 (pRS406-ADH1-NLS-Q2-mCherry) was generated by amplifying the NLS-Q2-mCherry from pRS313-CYC1-NLS-Q2-mCherry (Gift from A. Köhler) using Q5 polymerase. The PCR product was subsequently gel purified and assembled using Gibson Assembly (New England BioLabs) into pSR406-ADH1 digested with *Eco*RI*/Xba*I (New England BioLabs).

For generation of pDEST15-Chm7, used for bacterial expression of GST-Chm7 in lipid membrane strip binding experiments, the YJL049W cDNA coding for Chm7 (clone ScCD00011477) was supplied in pDONR201 by the Plasmid repository at Harvard Medical School ((Hu et al., 2007); https://plasmid.med.harvard.edu/) The construct was shuttled into the pDEST15 gateway plasmid (Thermo Fisher) carrying an N-terminal GST tag for bacterial protein expression and purification.

### Plate growth assays

To test for synthetic genetic interactions between *CHM7* and *PAH1* (DTCPL1634, 1659-60) or for growth complementation of *chm7Δapq12Δ* strains (PCCPL249, CPL1323, DTCPL1838, DTCPL1884-85, DTCPL1888, DTCPL1997-98), cells were cultured overnight at the permissive temperature. Cultures were diluted to an OD_600_ of 0.4 and spotted onto YPD in tenfold serial dilutions. Plates were wrapped in parafilm and incubated at either 23°C, 30°C, or 37°C for 36-48 h before imaging.

### Microscopy

To assess the localization of GFP fusions under the control of the *GAL1* promoter, strains (DTCPL515-16, DTCPL2030, DTCPL2056-60, DTCPL2066, DTCPL67) were grown in YPAR to mid-log phase. Expression of GFP fusions was induced upon addition of 1% galactose for 90 min prior to imaging.

To visualize the localization of intracellular PA and DAG, chm7_OPEN_-GFP-expressing cells (DTCPL413) containing pRS313-CYC1-NLS_Nup60_-Q2-mCherry or pRS313-CYC1-NLS_Nup60_-C1a-C1b-mCherry (Romanauska and Köhler, 2018) were cultured overnight at 30°C in SC-histidine, diluted into 10 ml of SC-histidine and cultured for at least 5 h before imaging.

For imaging the localization of cells expressing Chm7-GFP under its endogenous promoter (DTCPL81, DTCPL1861), strains were grown in YPAD.

To visualize NLS-PA sensor-mCherry in temperature-shift experiments in the *apq12Δ* and *nup116Δ* strains (DTCPL1979, DTCPL2035, DTCPL2039, DTCPL2050), cells were grown at the permissive temperature and shifted to 37°C for 3 h before imaging. Similarly, for colocalization of NPCs and the PA sensor, cells expressing GFP-Nup49 from pUN100-GFP-Nup49 (Belgareh and Doye, 1997) were grown in SC-leucine at 23°C and then shifted to 37°C for 3 h.

Cells were prepared for imaging as follows: Mid-log phase cells were collected by centrifugation at 375 g in a benchtop centrifuge for 1 min. Cells were resuspended in SC + 2% glucose and imaged immediately directly on the cover glass. Images were acquired on an Applied Precision DeltaVision microscope (GE Healthcare), using a 100x 1.4 NA oil immersion objective (Olympus) and a CoolSnapHQ^2^ CCD camera (Photometrics).

### Image processing/analysis

All micrographs presented were deconvolved using an iterative algorithm in softWoRx (6.5.1; Applied Precision GE Healthcare). Micrographs and gel images were further processed in FIJI/ImageJ (Schindelin et al., 2012) and Adobe Photoshop. Deconvolved images were used for line profiles and colocalization analyses, but for all quantification of fluorescence intensity unprocessed images were used.

To calculate a nuclear to cytosolic ratio of the NLS-GFP reporter (PLPC18), the average fluorescence of regions of the cytoplasm and nucleus within a middle z-section of individual cells was measured and related.

### Leptomycin B treatment

To inhibit nuclear export, experiments were performed as previously described (Thaller et al. 2019). In brief, KWY175 (gift from B. Monpetit) expressing GFP-fusions of Chm7 (DTCPL1815, DTCPL2051-2053), were grown to mid-log phase in YPAR at 30°C. 20% galactose was added to the culture medium to a final concentration of 1%. After 45 min, glucose (final [2%]) was added to the culture medium to arrest expression of GFP. 2 mL of the culture was then treated with either 5 µl of MeOH or LMB dissolved in 7:3 MeOH:H_2_O solution (Roche) at a final concentration of 50 ng/mL for 45 min before imaging.

### Repression of herniation formation in nup116Δ

To rescue the temperature sensitive herniation appearance in *nup116Δ* cells with *BRL1* overexpression, *nup116Δ* cells expressing the NLS-PA sensor-mCherry sensor (DTCPL2050) and harboring either pRS425 (2 μm/*LEU2*) or pGP564 (Hvorecny and Prelich, 2010) were grown at 23°C in SC-leucine (Sunrise Science). Cultures were split and kept at 23°C or shifted to 37°C for 3 h before imaging.

### Statistical methods

All graphs were generated using Prism (GraphPad 8.4). Appropriate statistical significance tests were selected as indicated in the figure legend and all values were calculated within Prism (GraphPad 8.4). *p*-values are indicated on the graph or in figure legends as: *ns, p* > 0.5; * *p* ≤ 0.05; ** *p* ≤ 0.01; *** *p* ≤ 0.001; **** *p* ≤ 0.0001. Error bars indicate the standard deviation from the mean. To measure the colocalization of GFP-Nup49 and the PA sensor at the nuclear envelope, the integrated density of GFP-Nup49 and NLS-PA sensor-mCherry were measured from a freehand 4 pixel-wide line profile drawn around the nuclear envelope of 15 randomly selected cells and measured using the Plot Profile plugin in FIJI (Schindelin et al., 2012). Corresponding values were plotted as a scatterplot and the linear correlation coefficient (Pearson’s, *r*) was calculated in Prism (GraphPad 8.4).

### Recombinant GST fusion protein production and purification

To produce recombinant fusion proteins, *E. coli* BL21 containing expression plasmids encoding for GST, GST-Chm7, GST-chm7-(W3AV1A), and GST-chm7-N, were grown in Terrific Broth (TB, 2.4% bacto-yeast extract (BD), 1.2% Bacto-tryptone (BD), 0.4% glycerol (Sigma)) overnight at 37°C, diluted to an OD_600_ of 0.1 and grown at 37°C until reaching an OD_600_ between 0.6-0.8. Subsequently, IPTG was added to a final concentration of 0.5 mM and cells were cultured at 23°C for 6 h. The bacteria were collected by centrifugation for 10 min at 10,000 rpm at 4°C, washed with ice-cold water and pelleted again at 4,000 rpm for 10 min at 4°C. Cell pellets were stored at -80°C.

Frozen cell pellets were resuspended in 15 ml of lysis buffer (50 mM Tris pH 7.4, 150 mM NaCl, 2 mM MgCl_2_, 10% glycerol) with the addition of 150 mM NaCl, 1 mM DTT, Roche protease inhibitors and 10 mg of lysozyme. The suspension was incubated on ice for 5-10 min and then lysed by sonication (Branson Sonifier 450). Lysates were pelleted at 11,700 rpm for 20 min at 4°C to remove cell debris. The supernatant was collected and incubated with 450 µL of glutathione sepharose 4B (GE Healthcare) for 1 h at 4°C. Beads were then collected by centrifugation and washed three times in lysis buffer. GST-fusions proteins were bound to the beads were eluted upon the addition of glutathione (10 mM in 50 mM Tris buffer pH 8.0) and the subsequent mixture was incubated in a rotor at 4°C for 20 min. Beads were pelleted and the supernatant was loaded into dialysis cassettes (3.5K, Slide-A-Lyzer, Thermo Scientific) that were then suspended in lysis buffer and dialyzed overnight to remove glutathione.

### Protein binding to lipid strip membranes

GST-Chm7 binding to lipid strips was performed following the recommended manufacturer protocol (Membrane Lipid Strips P-6002, Echelon Biosciences Inc.). Briefly, membrane lipid strips with 100 pmol of 15 different membrane lipids were incubated in blocking buffer (PBS-T: 50 mM Na_2_HPO_4_, 150 mM NaCl, pH 7.4, 0.1 % Tween-20, 3 % BSA fatty acid-free and globulin-free) overnight at 4°C. Recombinantly produced GST-Chm7 or GST only were added to a membrane at a final concentration of ∼68 µg/ml in PBS-T and incubated overnight at 4°C. After discarding the protein solution, the strips were washed 3 times in PBS-T before incubation in primary antibody (GST Tag Mouse anti-Tag, Clone: 8-326, Invitrogen) for 1 h at room temperature. The membrane was washed 3 times in PBS-T to remove any unbound primary antibody before incubating with the secondary antibody (anti-mouse IgG, HRP-linked antibody #7076 CST) for 1 h at room temperature. The membranes were washed in PBS-T before chemiluminescence detection.

### Liposome generation

Liposomes were prepared as follows: Glycerophospholipids dissolved in chloroform of POPA (1-palmitoyl-2-oleoyl-*sn*-glycero-3-phosphate; 840857C), POPI (1-palmitoyl-2-oleoyl-*sn*-glycero-3-phosphoinositol; 850142C), POPC (1-oleoyl-2-palmitoyl-*sn*-glycero-3-phosphocholine; 850457C), POPE (1-palmitoyl-2-oleoyl-*sn*-glycero-3-phosphoethanolamine; 850757C), were purchased from Avanti Polar Lipids. Lipid solutions were mixed in the desired ratios in a glass tube and were dried with N_2_ under a vacuum for 1 h. Dried lipid mixes were resuspended in SN buffer (20 mM Tris pH 7.6, 100 mM NaCl, 5 mM MgCl_2_) to a final concentration of 10 mM. The suspensions were transferred to Eppendorf tubes and subjected to 7 freeze/thaw cycles between liquid nitrogen and a 50°C water bath. Lipid mixtures were extruded 21 times through a double membrane filter paper with pores of the desired liposome diameter (50 nm, 200 nm, 400 nm). The collected liposomes were stored on ice at 4°C until use in binding/floatation assays. 25 nm liposomes were generated by extrusion through a 100 nm membrane followed by 3 consecutive 1 min. sonications (Branson Sonifier 450).

### Liposome binding and floatation assays

2 mM of prepared liposomes and 0.4 µM of selected GST fusions were mixed with SN buffer in a total volume of 150 µl in an Beckman 5×41 mm ultracentrifuge tube at 30°C for 1 h. Samples were adjusted to 40% Nycodenz by gently mixing with 150 µl of 80% Nycodenz in SN buffer. The mixture was layered with 250 µl of 30% Nycodenz in SN buffer and a top layer of 50 µl SN buffer. Samples were spun at 48,000 rpm at 4°C for 4 h in a Beckman SW Ti swinging-bucket rotor. 60 µL of floated sample (top layer) was collected using low retention pipet tips. Laemmli SDS-PAGE sample buffer was added, and samples were denatured at 95°C for 5 min. Proteins were resolved on an SDS-PAGE gradient gel (4-20%; BioRad) and visualized by SimplyBlue SafeStain (Thermo Fisher).

